# Validating indicators of CNS disorders in a swine model of neurological disease

**DOI:** 10.1101/514398

**Authors:** Vicki J. Swier, Katherine A. White, David K. Meyerholz, Aude Chefdeville, Rajesh Khanna, Jessica C. Sieren, Dawn E. Quelle, Jill M. Weimer

## Abstract

Genetically modified porcine disease models are becoming increasingly important for studying molecular, physiological and pathological characteristics of human disorders. Given their limited history, there remains a great need for proven reagents in swine tissue. To provide a resource for neurological models of disease, we validated antibodies by immunohistochemistry for use in examining central nervous system (CNS) markers. To validate these tools in a relevant model, we utilized a recently developed miniswine model of neurofibromatosis type 1 (NF1). NF1 is a tumor predisposition disorder, presenting with different type of tumors. Additionally, neurological associated symptomologies may include chronic pain, cognitive impairment, and behavioral abnormalities, making this miniswine model an ideal candidate for validating CNS-relevant antibodies. We validate antibodies implicated in glial inflammation (CD68), oligodendrocyte development (NG2, O4, Olig2, and myelin PLP), and neuron differentiation and neurotransmission (doublecortin, GAD67, and tyrosine hydroxylase) by examining cellular localization and brain region specificity. Additionally, we confirm the utility of anti-GFAP, anti-Iba1, and anti-MBP antibodies, previously validated in swine, by testing their immunoreactivity across multiple brain regions in mutant *NF1* samples. These validated immunostaining protocols for CNS markers provide a useful resource, furthering the utility of the genetically modified miniswine for translational and clinical applications.

## Introduction

Animal models are essential tools for studying the underlying mechanisms of disease as well as providing a platform for preclinical research and drug discovery. Historically, rodents have been one of the primary model systems for studying disease and driving drug discovery, largely due to the widespread availability of well-described and validated reagents for use in these model organisms. However, there are increasing instances where rodent models either fail to recapitulate aspects of human disease, or that treatments that are efficacious in a rodent model fail to translate to viable human therapies. This has led to development of large animal models of disease, such as genetically modified swine, which are more similar to humans anatomically, genetically, physiologically, and metabolically.^1-4^ This increased similarity is especially important when studying neurological disorders, where the anatomical and physiological differences found in the rodent systems, cannot recapitulate human disease. Successful genetically modified miniswine models have been established to study a number of human diseases including atherosclerosis, cancer, ataxia telangiectasia, cystic fibrosis, and neurofibromatosis type 1.^1,4-7^ However, while these new larger animal systems can better recapitulate many of the hallmarks of human disease, there are limited tools and reagents that have been well described and validated in these models. Herein, we test a number of antibodies relevant to the study of the brain and neurological disorders in porcine brain tissue.

We focus on reagents specific to neurons and glia (astrocytes, microglia, and oligodendrocytes) as these cells function together to support and protect neurons. When this function is disrupted, glia are implicated as causal agents in neurological disease, including Alzheimer’s, Huntington’s, and Parkinson’s disease.^8,9^ For example, dysregulation of oligodendrocytes, the myelinating cells of the CNS, can lead to loss of myelination. In other cases, a decrease in the density of oligodendrocytes in the prefrontal cortex may lead to schizophrenia, bipolar disorder, and major depressive disorder.^10^ Astrocytes, the most abundant cell-type in the CNS, serve as a neuronal support cell to promote survival,synaptogenesis and synapse pruning. In response to injury, astrocytes proliferate and/or are “activated” [indicated pathologically by an upregulation of glial fibrillary acidic protein (GFAP)] during various neurodegenerative diseases such as ALS and Parkinson’s disease.^8^ Microglia, the primary immune cells in the CNS, can sense changes in their environment and either promote healthy neurons or provide protection to neurons that have been injured or diseased.^11^ Microglia produce proinflammatory agents that recruit inflammatory cells that are toxic to neurons, contributing to neurodegenerative diseases like multiple sclerosis.

Neurons are highly involved in signal transmission within the CNS.^12^ They release chemical neurotransmitters that affect signaling between neurons and play a role in various physiological functions of the CNS. A loss of neurons in specific regions of the CNS may cause certain affects, for example, loss of dopaminergic neurons in the substantia nigra in patients with Parkinson’s disease, causes reduced balance and motor coordination.^13^

Several studies of swine models of CNS injuries/disorders have tested of the specificity of antibodies within the CNS.^14-18^ However, two of these studies have focused solely on the spinal cord and not the brain itself.^15,16^ Of the remaining studies, only one of these describes the impact of CNS injuries/disorders by addressing axonal injury and astrocytic/microglial reactivity.^14^ The other brain immunohistological study was similar to ours as it validated antibodies, however this validation was compared to a general histological stain (Giemsa).^17^ Moreover, we recently published a comprehensive study on antibody immunoreactivity in swine tissues, specifically wild type swine,^19^ but more studies are needed to validate markers that have a role in CNS disorders, specifically in genetically modified swine that recapitulate characteristics of a human disease. Anti-GFAP and anti-Iba1 antibodies have been used in a number of swine studies, however, none have explored expression/activation within specific brain regions (that may be impacted due to disease).

As some antibodies are predominantly reactive in a disease state, here we use a recently developed miniswine model of NF1 to validate a number of CNS cell-specific antibodies.^1^ As patients with NF1 experience a host of CNS-specific impairments, this porcine model is ideal for validation of neurologically relevant antibodies. We validate and explore the expression of antibodies implicated in glial inflammation, oligodendrocyte differentiation, neuronal signaling, and nociceptive function. Taken together, we provide a powerful set of tools to researchers modeling neurological dysfunction in porcine models of disease.

## Materials and Methods

### Animal Tissue

All miniswine were maintained at Exemplar Genetics under an approved IACUC protocol. All mice were maintained in an AAALAC accredited facility in strict accordance with National Institutes of Health guidelines, and studies were approved by the Sanford Institutional Animal Care and Use Committee (USDA License 46-R-0009).

### Tissue Microarray

Regions from formalin fixed cortex (CTX), cerebellum (CB), hippocampus (HPC), thalamus (THAL), corpus callosum (CC), and cerebral aqueduct (CGG) of a 15-month old, male *NF1* miniswine^1^ were isolated and placed in tissue cassettes. These regions were selected due to their relevance to neurologic disease in relation to macrocephaly (CC),^20^ white matter abnormalities (CTX, CB, and CC)^21^ brain lesions (CB and THAL),^22^ abnormal physiology (HPC),^23^ and aqueductal stenosis (CGG).^24^ The tissue cassettes were processed in a Lecia ASP300 Tissue Processor (Lecia Biosystems Inc, Buffalo Grove, IL) and embedded in paraffin. Sections from each paraffin block were cut with a Leica RM2125 (Lecia Biosystems Inc, Buffalo Grove, IL). Subsections of interest were marked on each slide and a circular biopsy was taken from the paraffin block that matched the marked region. The paraffin biopsies were placed into a tissue microarray mold and re-embedded in paraffin to create a paraffin microarray block. Sections of the paraffin microarray block were cut and floated onto slides.

### Validation of antibodies

Details regarding each of the antibodies used in this study are listed in **Table 1**. When possible, we selected antibodies predicted to work in swine or constructed with a porcine immunogen, however, very few of these antibodies exist. Therefore, we primarily selected focused on antibodies known to react in multiple mammalian species (such as mouse, rat and human), as the degree of homology between swine and the aforementioned mammalian proteins is fairly high (89-100% similarity),^25^ especially for evolutionarily conserved genes. More consistent immunopositive results in swine were obtained by selecting antibodies in this manner instead of antibodies that only react in human or mouse tissue.

**Table 1.**
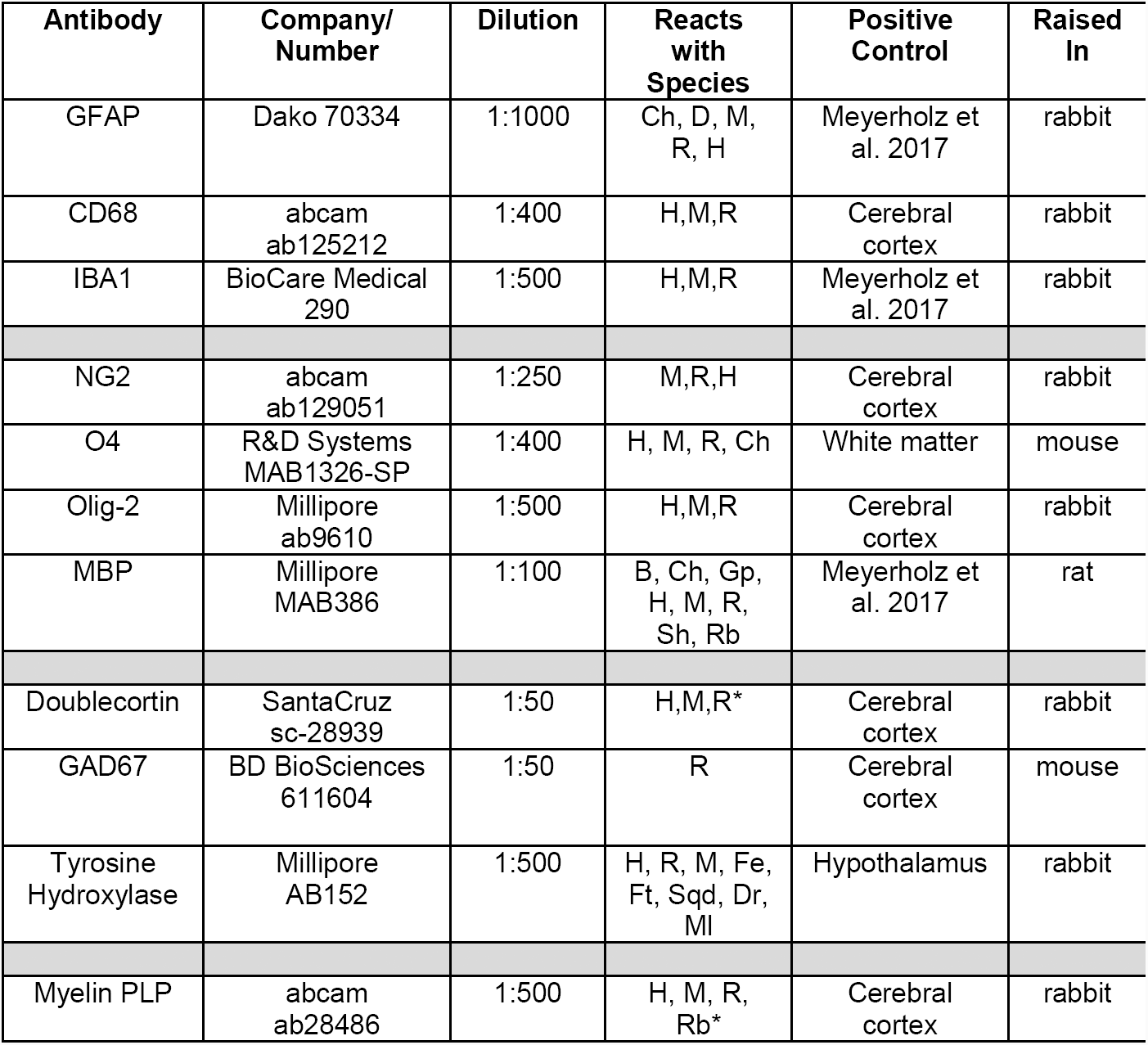
Description of antibodies used and dilutions. Antibody, company, catalog number, reaction with species, the host animal, target of the antibody, and cellular localization of each antibody examined. Target and cellular location were either cited from the antibody data sheet (provided by the company) or obtained from the Human protein atlas, https://www.proteinatlas.org/. * indicates predicted to react in swine.

The antibodies that we validated were known to react in postnatal mouse tissues based on information provided from the manufacturer. According to the gene expression database at the mouse genome informatics website (http://www.informatics.jax.org),^26^ the markers that we validated were known to have expression in the forebrain of the mouse (NG2), cerebral cortex and hippocampus (doublecortin, GAD67), and corpus callosum and hippocampus (myelin PLP). Other studies have found CD68 (macrosialin) expression in the corpus callosum and striatum of C57BL/6 mice,^27^ Olig-2 expression in corpus callosum and ventral forebrain regions,^28^ O4 protein expression in cerebral cortex and above the cingulum,^29^ and tyrosine hydroxylase protein expression in the forebrain-cerebral cortex and hippocampus of mice.^30^ As a positive control, a coronal section of mouse brain was immunolabeled alongside the miniswine tissue, to verify the proper reactivity, localization, and expression of the antibody in question.

Immunogen peptides sequences were obtained from manufacturer’s documentation and compared to swine (*Sus scrofa*) protein sequences from the Refseq database using the Basic Local Alignment Search Tool (BLAST) from NIH (**Table 2**).

**Table 2:**
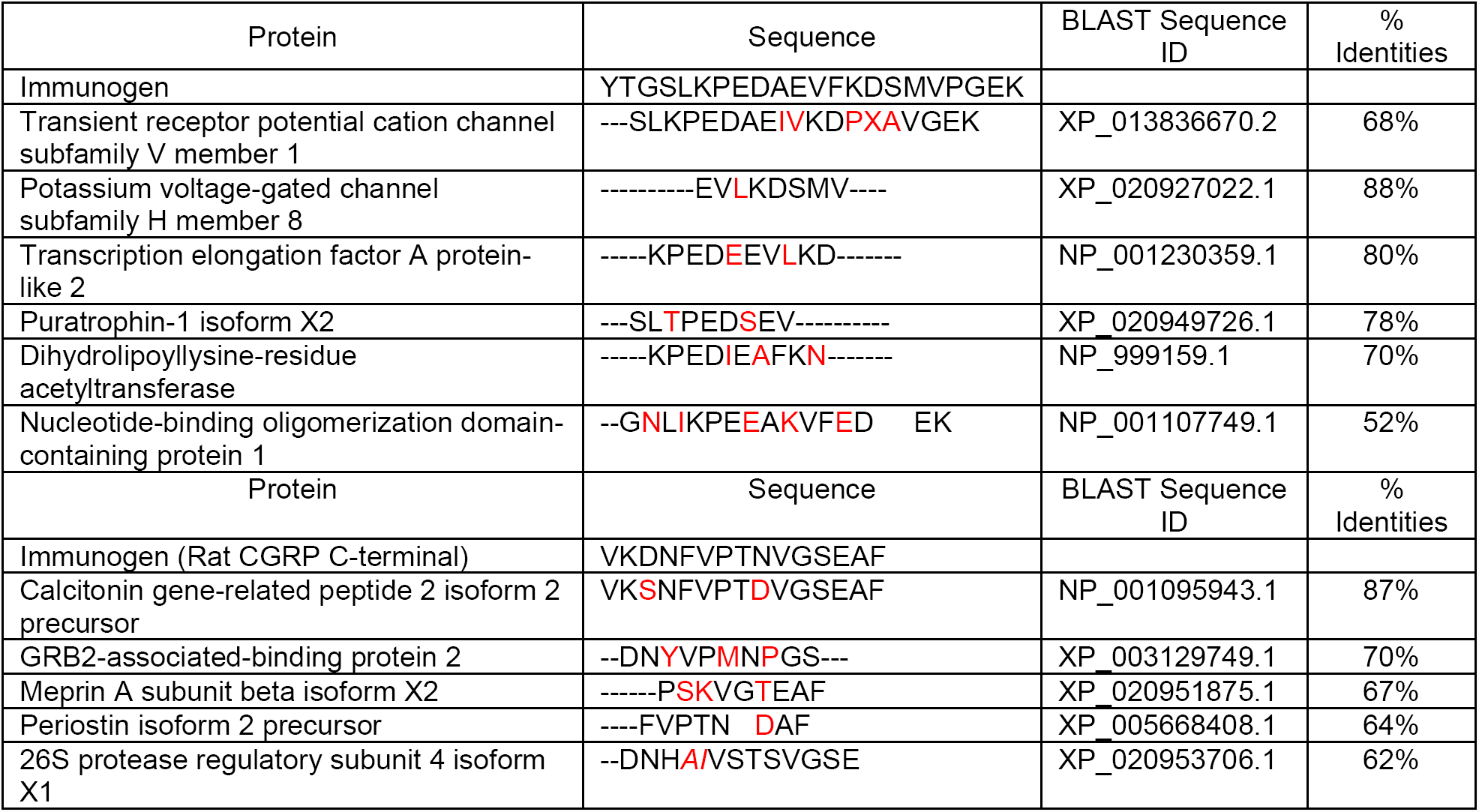
Sequence identities between immunogen peptides. used to generate the commercial antibodies against TRPV1 and CGRP and off-target pig proteins extracted from the Refseq database using BLAST. Amino acids differing from the immunogen peptides are represented in red and additional amino acids are in italic.

### Immunohistochemistry

Paraffin tissue arrays on slides were deparaffinized in xylene, rehydrated in ethanol, and rinsed in double distilled water. Antigen retrieval was performed at 90C for 20 minutes using sodium citrate buffer, pH 6. Then, slides were rinsed in 1xTBST, endogenous peroxidases were blocked in Bloxall™ (Vector Laboratories, Burlingame, CA) for 10 minutes, and rinsed again in 1xTBST. *<u>For antibodies raised in rabbit</u>*, blocking serum from an ImmPRESS™ HRP Anti-Rabbit IgG (Peroxidase) Polymer Detection Kit (Vector Laboratories) was incubated on slides for 20 minutes at room temperature (RT). Slides were drained and the primary antibody was incubated on the slides at 4C overnight. Negative controls without primary antibody were run in parallel with normal host IgG. Slides were rinsed in 1xTBST, and ImmPRESS™ (Peroxidase) Polymer was incubated on slides for 30 minutes at RT. Slides were rinsed in 1xTBST and 3,3’-diaminobenzidine (DAB) from Vector laboratories was added to the slides for 2 to 10 minutes until the DAB activation occured. Slides were then washed with DI water, stained with Mayer’s hematoxylin, washed with running tap water, dipped in 0.25% Lithium carbonate, rinsed in DI water, dehydrated with ethanol, cleared with Xylene and mounted with DPX mounting media. *<u>For antibodies raised in mouse</u>*, an ImmPRESS™ HRP Anti-Mouse IgG (Peroxidase) Polymer Detection Kit (Vector Laboratories) was used, followed by the described DAB staining. *<u>Forantibodies raised in rat,</u>* 1% goat serum with Triton was used as a blocking solution for at least 1 hour at RT and a Goat Anti-Rat IgG H&L (HRP) (Abcam ab97057) was used as a secondary and incubated for 1.5 hours at RT before continuing with the described DAB staining. Tissue sections were viewed with an Aperio Versa slide scanner (Lecia Biosystems Inc, Buffalo Grove, IL) and images were extracted with Leica’s ImageScope software.

### Dorsal root ganglia (DRG) Immunostaining

Miniswine DRGs were fixed in 4% PFA for 24 hours, cryoprotected in 30% sucrose (m/v) in PBS for 48 hours and frozen at -70°C in 2-methylbutane chilled with dry ice. Samples were cut into 20µm-thick sections. Sections were subsequently blocked in 3% BSA, 0.1% Triton X-100 in PBS for 1 hour at room temperature then incubated with primary antibodies (anti-TRPV1, Neuromics GP14100; anti-CGRP, Abcam ab16001) diluted in blocking solution overnight at +4°C. After 3 washes in PBS, slides were incubated with secondary antibody diluted at 1/1000 in blocking solution for 2 hours at room temperature, washed and counterstained with DAPI. For the negative control, primary antibodies were omitted. Images were acquired on an Axio Imager 2 (Zeiss), using a 10X objective controlled by the Zen software (Zeiss).

## Results

As research on CNS-related antibodies for immunohistochemistry are lacking in swine models of human disorders, we validated glia and neuron related antibodies in a model of NF1.

### Immunolabeling of known markers in various regions of *NF1* mutant miniswine brain

GFAP, a marker of intermediate filaments in astrocytes that become hypertrophic in response to insult, has been shown to be increased in a number of neurological diseases, including NF1.^31^ Here, we observed classic star-shaped GFAP^+^ immunostaining in a 15-month-old *NF1* mutant miniswine (Fig. 1A-C, arrows). The cytoplasmic localization pattern was similar to that seen previously in human and swine cerebellum.^19^ Immunopositive cells were present in the *NF1* miniswine cortex, cerebellum and thalamus, mirroring what has been observed in identical regions of the human brain.^32^

**Figure 1:**
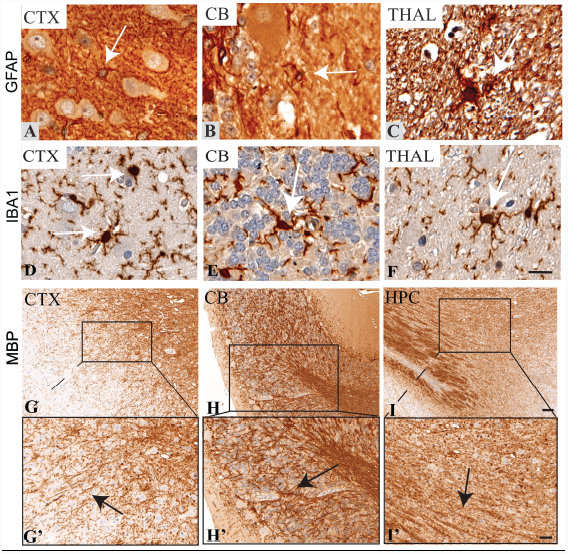
Immunolabeling of known markers in various regions of *NF1* mutant miniswine brain. A-C: GFAP+ expression in the cortex (CTX), cerebellum (CB), and thalamus (THAL). Arrows indicate classic, star-shaped morphology of astrocytes. D-F: Iba1+ expression in the cortex, cerebellum, and thalamus. Arrows indicate both ramified and ameboid-like microglia. Scale bar: 50µm. G-I: MBP+ expression in the cortex (CTX), cerebellum (CB), and hippocampus (HPC). Scale bar: 500µm. G’-I’: Enhanced magnification of panels A-C. Scale bar: 200µm.

Cytoplasmic ionized calcium binding adaptor molecule 1 (Iba1) immunostaining was observed in both non-reactive, ramified cells with numerous branching processes and more reactive amoeboid-like cells (arrows) in the cortex, cerebellum and thalamus (Fig. 1D-F). Filamentous immunopositive structures likely represent cross-sectional microglial processes. The cytoplasmic localization is similar to published results in human and swine cerebrum (also referred to as AIf1).^19^ We expected to document Iba1 staining in various regions of the mutant miniswine brain, as injury and inflammatory factors activate Iba1+ microglia in affected areas of the brain, such as the cerebellum of sheep exposed to LPS,^33^ and thalamus of wild type mice exposed to traumatic brain injury.^34^

Large areas within the cortex, cerebellum and hippocampus were immunopositive for anti-myelin basic protein (MBP) antibodies, which label mature oligodendrocytes in white matter tracts (Fig. 1G-I, G’-I’, arrows). A higher magnified image of the tracts documents the filamentous morphology of the cytoplasmic and membranous localization of MBP to the myelin sheath (Fig. 1G’-I’). We see a similar localization pattern to the results shown in cerebral white matter of humans and swine.^19^ The results are consistent with the expectation of MBP immunostaining in any white matter tracts of the brain, which include regions of the cortex, cerebellum, and hippocampus.

### Detection of microglia and oligodendrocyte cell lineage in the *NF1* miniswine brain

Reactive microglia, indicated by cytoplasmic CD68^+^ immunostaining, were localized to the cytoplasm (arrows) in the cortex, cerebellum, and hippocampus (Fig. 2A-C). We observed a similar cytoplasm localization in mouse cerebral cortex (Fig. 2D). CD68 immunoreactivity has been documented in microglia within the cerebral cortex of humans,^32^ and in the multiple brain regions of aging wild type mice and mice exposed to an neurological insult such as LPS.^35^

**Figure 2:**
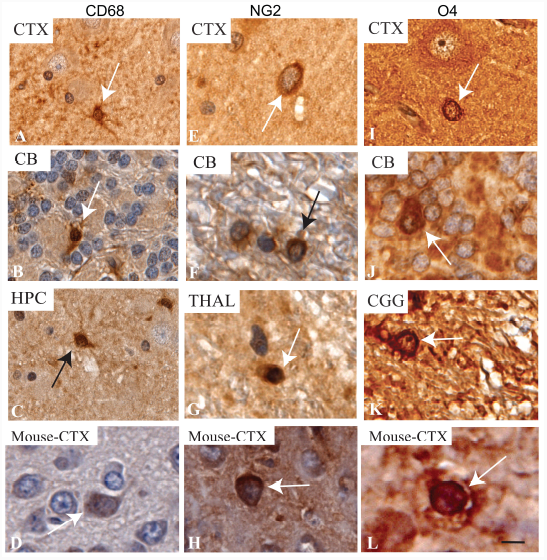
Immunolabeling of microglial and Pre-oligodendrocyte markers in the *NF1* mutant miniswine brain. A-C: CD68^+^ expression in the cortex, cerebellum, and hippocampus (HPC). Arrows indicate ameboid-like microglia. D: CD68+ expression in mouse cerebral cortex. E-G: NG2+ expression in the cortex (CTX), cerebellum (CB), and thalamus (THAL). Arrows indicate immature oligodendrocytes. H: NG2+ expression in mouse cerebral cortex. I-K: O4+ expression in the cortex, cerebellum, and cerebral aqueduct (CGG). Arrows indicate pro-oligodendrocytes. L: O4+ expression in mouse cerebral cortex. Scale bar: 50µm.

Tracing cell type lineage can be incredibly informative in determining mechanisms of disease and appropriate intervention points. Oligodendrocytes have a well-studied lineage and are of particularly important for the pathology of NF1. For example, oligodendrocyte precursor cells (OPCs) have been identified as the cell of origin for gliomas in *NF1* conditional knockout mice^36^, with *NF1* mutant mice expressing more OPCs in the brain. As expected, we observed membrane and cytoplasmic neural/glial antigen 2+ (NG2) immunostaining in the cortex, cerebellum and thalamus, indicating the presence of oligodendrocyte progenitors (Fig. 2E-G, arrows). We see a similar membrane and cytoplasmic localization in mouse cerebral cortex (Fig. 2H). NG2+ immunostaining has been documented in the cerebral cortex and cerebellum in human tissues,^32^ and throughout the rodent brain, including the thalamus of wild-type rats.^37^ Oligodendrocyte marker 4+ (O4) immunostaining to the membrane indicates the presence of pre-oligodendrocytes in the cortex, cerebellum, and cerebral aqueduct (Fig. 2I-K, arrows). Compared to mouse cerebral cortex, we see a similar membrane localization pattern (Fig. 2L). O4+ immunostaining has been found in the corpus callosum in young rat pups,^38^ in the rat cerebellum,^39^ and in the midbrain (substantia nigra pars compacta) of control and neurotoxin exposed C57BL/6 mice.^40^

### Detection of matured oligodendrocytes in the *NF1* miniswine brain

The nucleus and cytoplasm of multiple oligodendrocytes (in various stages of differentiation) were immunopositive for anti-Olig2 antibodies in the cortex, cerebellum, and cerebral aqueduct (Fig. 3G-I, arrows). Comparatively, we see evidence of nuclear immunostaining in mouse cerebral cortex (Fig. 3D). Though Olig2 was shown to be expressed in all oligodendrocyte lineages^41^, it is known to traditionally label immature oligodendrocytes, thus is most abundant in the developing brain. In the adult human brain, Olig2^+^ cells can be found in the cerebral cortex, in molecular and granular layer cells of the cerebellum,^32^ and in similar regions in mice.^28,42^ As a marker of mature oligodendrocytes, we also tested a myelin proteolipid protein (Myelin PLP) antibody with inconsistent results. There was faint membrane expression in the white matter tracts of the cortex (Fig. 3E), cerebellum (Fig. 3F) and corpus callosum (Fig. 3G) to the neuropil, though there appears to be non-specific staining in other cells within the CNS. We see similar staining to the neuropil and non-specific immunostaining in mouse cerebral cortex (Fig. 3H).

**Figure 3:**
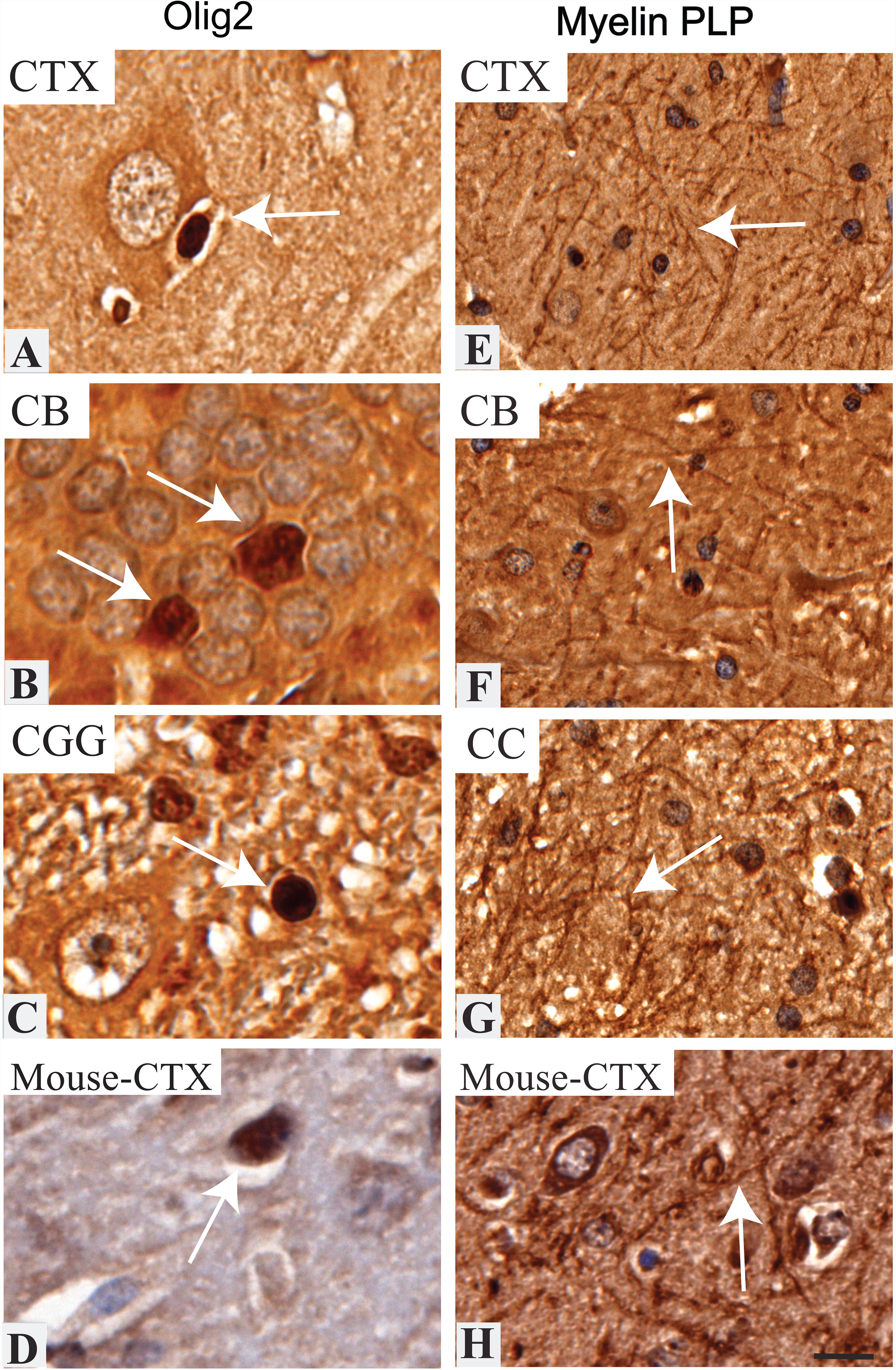
Immunolabeling of mature oligodendrocyte lineage markers in the *NF1* mutant miniswine brain. A-C: Olig2^+^ expression in the cortex, cerebellum, and cerebral aqueduct. Arrows indicate several oligodendrocytes in different developmental stages. D: Olig2+ expression in mouse cerebral cortex. E-G: Myelin PLP+ expression in the cortex (CTX), cerebellum (CB) and corpus callosum (CC). H: Myelin PLP+ expression in mouse cerebral cortex. Scale bar: 50µm.

### Immunolabeling of various neuronal subtypes and neurotransmitters in the *NF1* miniswine brain

Spatial learning and memory deficits are a prominent feature of neurological disease, specifically affecting dopamine, GABA, and glutamate signaling in hippocampal neurons.^23,43^ Dysregulated GABA signaling in the CNS, which causes an increase of GABA-mediated inhibition, has been implicated as a cause of learning defects in mice models of Huntington’s disease and NF1.^23,44^ Loss of dopamine signaling in hippocampal neurons has been proposed to be the reason for the spatial learning and memory defects in mutant *NF1* mutant mice and a treatment of dopamine to these mice rescued the long-term potentiation response to normal levels.^43^

In our study, we report cytoplasmic doublecortin (DCX)^+^ immunostaining in the cortex, cerebellum and hippocampus of the adult miniswine brain, indicating the presence of immature neurons (Fig. 4A-C). Cytoplasmic immunostaining of DCX was also found in mouse cerebral cortex (Fig. 4D). This was as expected, as DCX+ immunostaining has been documented in the cerebral cortex and hippocampus in mice,^26^ and in the granular cell layer of the adult rat cerebellum.^45^ Glutamate decarboxylase 67 (GAD67) immunopositive cells were found in the cortex, cerebellum, and hippocampus in the *NF1* miniswine brain (Fig. 4E-G), as expected, since GAD67+ neurons have been documented within these brain regions in humans.^32^ Inhibitory interneurons that express GAD67 are small and /or medium-sized oval shaped cells with cytoplasmic expression in the cell body and dendritic processes.^46^ Very clear immunostaining with anti-GAD67 antibodies was found in the cytoplasm of these neurons, specifically localized to the cell bodies (Fig. 4E-G; single arrow). We see similar cytoplasmic and membranous immunostaining of GAD67 in mouse cerebral cortex (Fig. 4H). Prominent cytoplasmic tyrosine hydroxylase (dopaminergic neurons) immunostaining was found in the cerebral aqueduct, positive immunostaining in the cortex and in the cerebellum (Fig. 4I-K). This immunostaining matches the cytoplasmic localization of tyrosine hydroxylase in mouse cerebral cortex (Fig. 4L). The immunoreactivity of tyrosine hydroxylase is as expected, as this marker has been found to immunoreact to the cerebellar lobules and Purkinje cells of the cerebellum in adult mice;^47^ and similarly in wild type rats, the somata of cortical tyrosine hydroxylase-immunoreactive interneurons were less than 15 µm in diameter and the cell bodies were primarily fusiform and bipolar shaped.^48^

**Figure 4:**
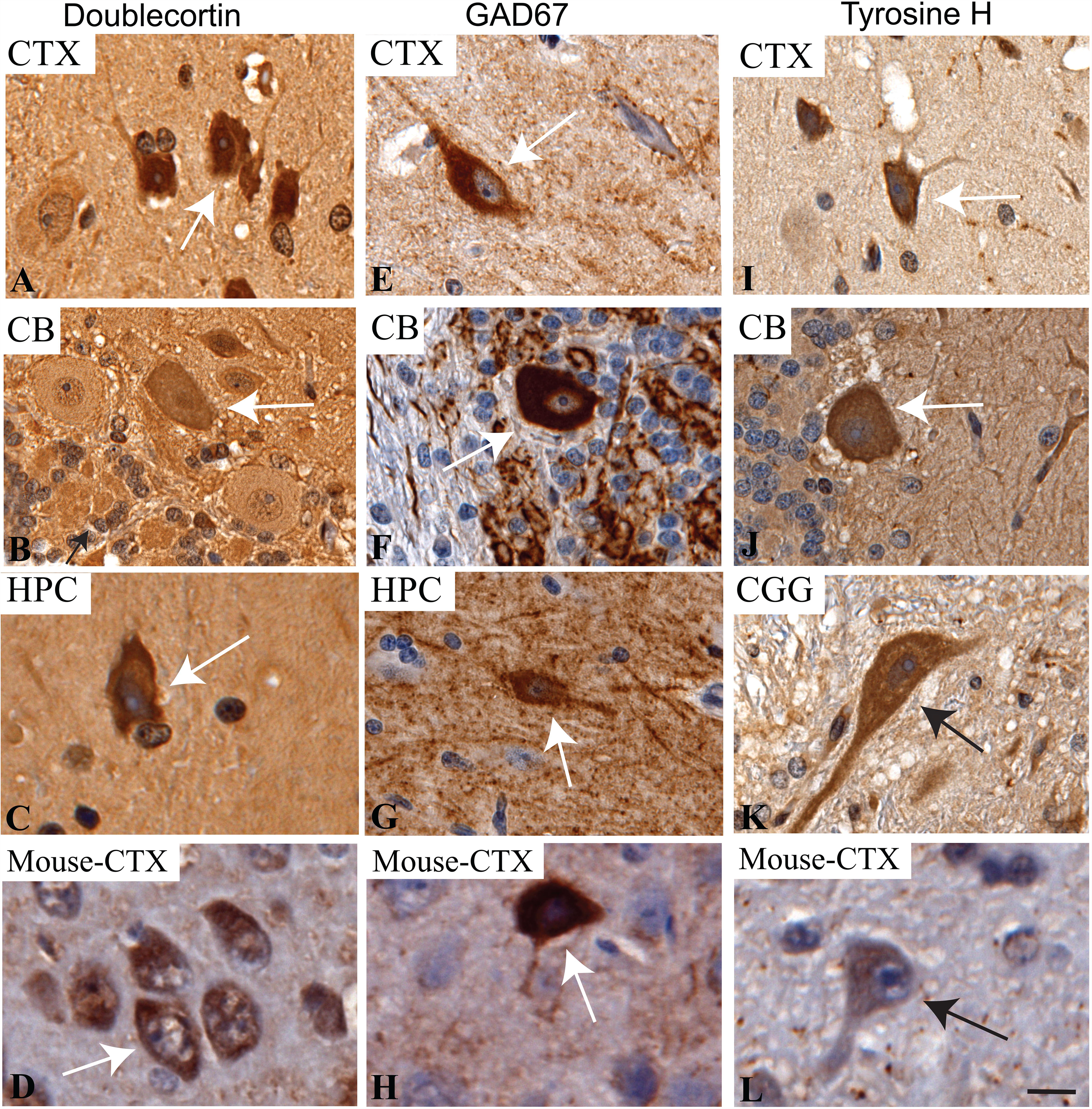
Immunolabeling of several neuron markers in the *NF1* mutant miniswine brain. A-C: Doublecortin^+^ expression in the cortex (CTX), cerebellum (CB), and hippocampus (HPC). Arrows indicate immature neurons, with minimal staining in the cerebellum. D: Doublecortin+ expression in mouse cerebral cortex. E-G: GAD67^+^ expression in the cortex, cerebellum, and hippocampus. Arrows indicate GABAergic neuron cell bodies. H: GAD67+ expression in mouse cerebral cortex. I-K: Tyrosine H^+^ expression in the cortex, cerebellum, and cerebral aqueduct (CGG). Arrows indicate dopaminergic neurons. L: Tyrosine H+ expression in mouse cerebral cortex. Scale bar: 50µm.

### Nociceptive markers tentatively identified in miniswine DRGs of the spinal cord

Chronic pain is usually a component of many neurological diseases that affects approximately 20-40% of patients.^49^ We investigated the utility of pain perception-nociceptive markers in dorsal root ganglion isolated from our mutant miniswine. Sliced DRGs from miniswine were immunostained with antibodies raised against calcitonin gene related peptide (CGRP), a pro-nociceptive neurotransmitter, and transient receptor potential cation channel subfamily V member 1 (TRPV1), a capsaicin receptor. Unlike immunostaining patterns observed in rodents and Rhesus monkeys, in which subpopulations of neurons were stained by each antibody; ^50,51^ all DRG neurons were stained by antibodies against CGRP and TRPV1 with different fluorescent intensities in miniswine (Fig. 5A-B). For one of the antibodies, differences in staining pattern were observed between batches. Control omitting primary antibodies revealed no non-specific staining due to the secondary antibodies (Fig. 5C-D). However, comparison of the immunogen sequences with swine (Sus scrofa) protein databases in **Table 2**, revealed that several off-target proteins could also bound by these antibodies. Therefore, better controls are necessary to validate these antibodies for use in swine.

**Figure 5:**
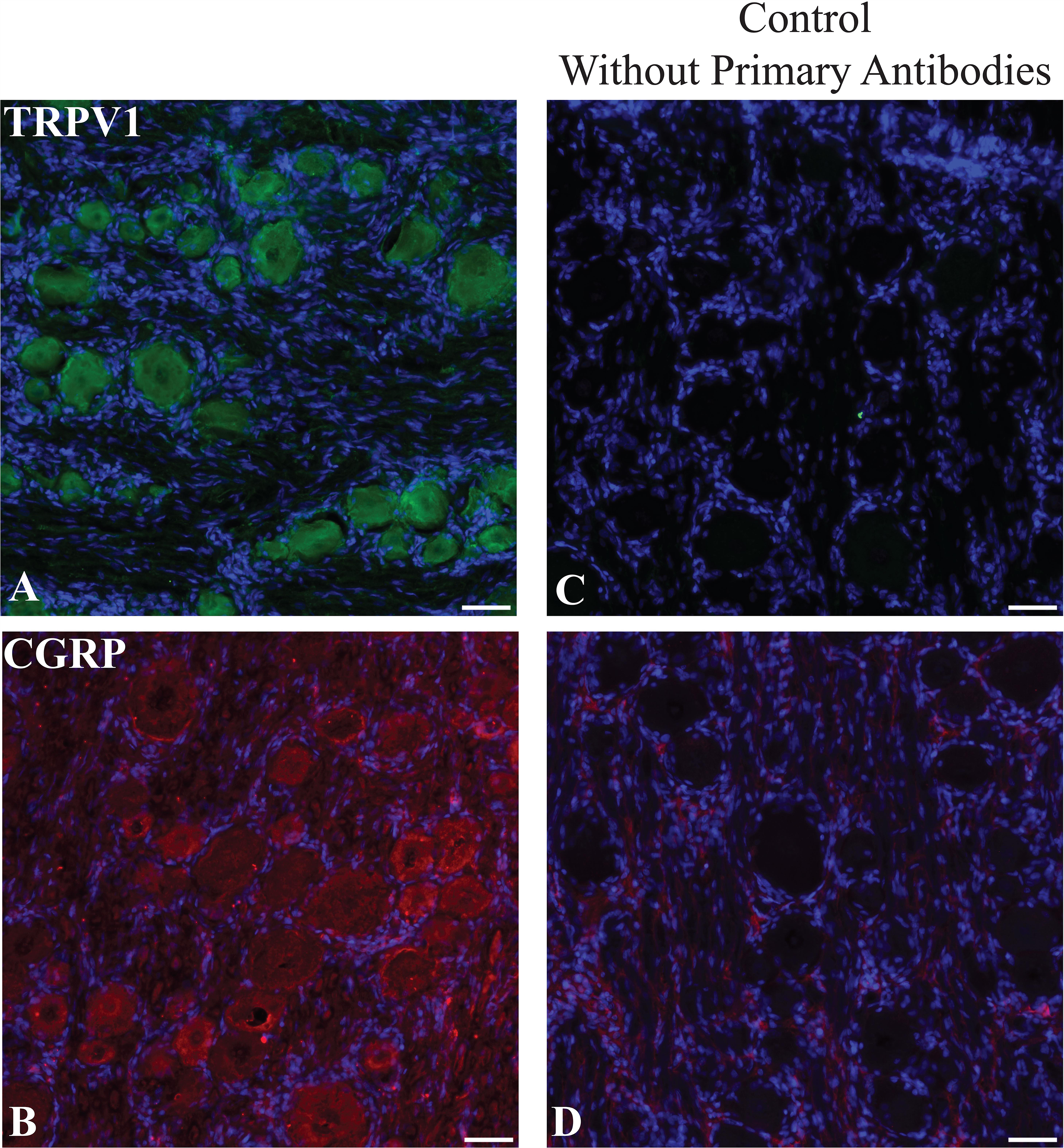
Immunolabeling miniswine DRGs. A-B: Miniswine DRGs were immunostained with commercial antibodies against TRPV1 and CGRP. C-D: Negative controls omitting primary antibodies. Nuclei were counterstained with DAPI. Scale bars: 50µm.

## Discussion

The paucity of known antibodies that are validated in swine creates roadblocks in experimental design in miniswine models of disease. In the field of neurological research, more translatable large animal models of diseases may be less likely to be constructed if appropriate reagents are not readily available. For example, while a particular antibody of interest may be available, they are more likely to react in humans or mice, requiring significant time and resources to optimize immunostaining conditions in porcine. Resources to test numerous antibodies is often limited and, ultimately, because of the lack of validated tools, it can place technical constraints on proper experimental design, making accessible resources such as this study critical in the swine model community.

Modeling a neurological disease may require the involvement of different types of CNS cells. Glia play an important role in the pathogenesis of neurological diseases, especially for documenting inflammation (Iba1+ and CD68+ microglia), astrocytosis (GFAP+ astrocytes), and disruption of myelination (NG2+ and O4+ OPCs, MBP and myelin PLP+ mature oligodendrocytes, Olig2+ all oligodendrocytes). These markers are widely studied as a hallmark of neurodegeneration as they commonly are found in early stages of neurological disease as an indicator that the microenvironment of the brain has been altered, and neuron function is likely to be disrupted. For example, upregulated Olig2 is an indicator that OPCs are activated in response to autoimmune regulated demyelination in multiple sclerosis.^52^ These markers continue to be pathological markers in later stages of neurological diseases such as Alzheimer’s. In postmortem Alzheimer’s patients, dense populations of CD68+ microglia exist in the hippocampus,^53^ and reduced NG2 immunoreactivity was found in brain tissue.^54^ Glial markers are very effective tools to document the progression of a disease. Studying their mechanisms of activation and differentiation during the pathogenesis of neurodegeneration will help to develop therapeutics that treat and prevent these diseases.

Neuronal inflammation, which can be caused by astrogliosis and activated microglia, can lead to disrupted neuronal transmission. Disrupted neuronal signaling and loss of neurons can lead to learning deficits, cognitive dysfunction, and loss of motor control. The neuronal markers that we studied, which are involved in GABA (GAD67+) and dopamine (tyrosine hydroxylase+) neurotransmission have benefits to neurological disease research that studies activity and function of neurons in conjunction with cognitive and motor impairment. For example, altered GABA level and synthesis is implicated in Huntington’s disease (HD), and transgenic mice that express exon 1 of the HD gene had reduced levels of GAD67 in brain tissues,^55^ and these mice have difficulty performing spatial cognition tasks.^44^

To study these dysfunctions in swine models, that provide more translatable human therapies, reagents specific to swine need to be validated. Ours is the first study validating antibodies specific for CD68, NG2, O4, Olig-2, GAD67, tyrosine hydroxylase, and myelin PLP in the CNS of mutant miniswine; and the first study validating GFAP, Iba1, MBP, and doublecortin in the cerebral cortex, cerebellum, thalamus, and hippocampus of miniswine. GFAP was validated in 100 day old swine in the neocortex.^17^ Our previous publication only validated GFAP, Iba1, and MBP in the cerebrum of wild type swine.^19^ Compared to our previous publication, we chose an anti-MBP antibody that was specific to a different amino acid chain (aa82-87 vs aa182-197), a different dilution (1:100 vs 1:600), and a different secondary (HRP conjugated secondary vs HRP Labeled Polymer). Our antigen retrieval process was longer for anti-Iba1 and anti-GFAP (20 minutes vs 5 minutes), the temperature of our antigen retrieval process was less (approximately 95C vs 110-125C), and we incubated our primary antibody overnight compared to 1 hour. We also chose a less concentrated dilution for anti-GFAP (1:1000 versus 1:200) and more concentrated for anti-Iba1 (1:500 vs 1:2000).

Though we were able to validate all these markers in various regions of a mutant miniswine brain, we only validated nociceptive markers in peripheral CNS tissues (i.e. spinal cord). Neurological diseases such as amyotrophic lateral sclerosis (ALS) have neuronal loss and degeneration in both the brain and spinal cord. Validating more CNS markers in the swine spinal cord would improve the utility of swine models that investigate neurodegeneration throughout the CNS. Moreover, we only validated markers of dopaminergic and GABAergic neurotransmission in this study, thus further work to validate markers specific for glutamatergic neurons (excitatory), serotonergic neurons (cognition, learning, memory), and cholinergic neurons (acetylcholine-motor) in miniswine models is still needed.

Validation of immunological tools (commonly tested on mice and human tissue) in swine tissues will improve the clinical and translational aspects of the swine model for disease research, The validation of the antibodies described in this paper provides new tools that will aid in the further investigation of the role of neurofibromin in cognitive dysfunction, as well as other swine model of CNS disease.

## Acknowledgements

We would like to acknowledge the following individuals for technical assistance: Brian Dacken, Trisha Smit, and Justin Van Kalsbeek of Exemplar Genetics. This work was supported by funding from the Children’s Tumor Foundation to DKM, RK, JCS, DEQ, and JMW. This work also received support from the Sanford Research Imaging Core (NIH P20GM103620) and Claire Evans in the Sanford Research Molecular Pathology Core (NIH P20GM103548).

## References

1. White KA, Swier VJ, Cain JT, et al. A porcine model of neurofibromatosis type 1 that mimics the human disease. JCI insight. 2018;3(12).

2. Walters EM, Wolf E, Whyte JJ, et al. Completion of the swine genome will simplify the production of swine as a large animal biomedical model. BMC medical genomics. 2012;5:55.

3. Prather RS, Lorson M, Ross JW, Whyte JJ, Walters E. Genetically engineered pig models for human diseases. Annual review of animal biosciences. 2013;1:203–219.

4. Rogers CS, Abraham WM, Brogden KA, et al. The porcine lung as a potential model for cystic fibrosis. American journal of physiology. Lung cellular and molecular physiology. 2008;295(2):L240–263.

5. Davis BT, Wang XJ, Rohret JA, et al. Targeted disruption of LDLR causes hypercholesterolemia and atherosclerosis in Yucatan miniature pigs. PloS one. 2014;9(4):e93457.

6. Sieren JC, Meyerholz DK, Wang XJ, et al. Development and translational imaging of a TP53 porcine tumorigenesis model. J Clin Invest. 2014;124(9):4052–4066.

7. Beraldi R, Chan CH, Rogers CS, et al. A novel porcine model of ataxia telangiectasia reproduces neurological features and motor deficits of human disease. Human molecular genetics. 2015.

8. Phatnani H, Maniatis T. Astrocytes in neurodegenerative disease. Cold Spring Harbor perspectives in biology. 2015;7(6).

9. Verkhratsky A, Parpura V. Neurological and psychiatric disorders as a neuroglial failure. Periodicum biologorum. 2014;116(2):115–124.

10. Vostrikov V, Uranova N. Age-related increase in the number of oligodendrocytes is dysregulated in schizophrenia and mood disorders. Schizophrenia research and treatment. 2011;2011:174689.

11. Hickman S, Izzy S, Sen P, Morsett L, El Khoury J. Microglia in neurodegeneration. Nature neuroscience. 2018;21(10):1359–1369.

12. Ransohoff RM. How neuroinflammation contributes to neurodegeneration. Science. 2016;353(6301):777–783.

13. Miller DW, Cookson MR, Dickson DW. Glial cell inclusions and the pathogenesis of neurodegenerative diseases. Neuron glia biology. 2004;1(1):13–21.

14. Kallakuri S, Desai A, Feng K, et al. Neuronal Injury and Glial Changes Are Hallmarks of Open Field Blast Exposure in Swine Frontal Lobe. PloS one. 2017;12(1):e0169239.

15. Navarro R, Juhas S, Keshavarzi S, et al. Chronic spinal compression model in minipigs: a systematic behavioral, qualitative, and quantitative neuropathological study. Journal of neurotrauma. 2012;29(3):499–513.

16. Raore B, Federici T, Taub J, et al. Cervical multilevel intraspinal stem cell therapy: assessment of surgical risks in Gottingen minipigs. Spine. 2011;36(3):E164–171.

17. Hou J, Riise J, Pakkenberg B. Application of immunohistochemistry in stereology for quantitative assessment of neural cell populations illustrated in the Gottingen minipig. PloS one. 2012;7(8):e43556.

18. Say M, Machaalani R, Waters KA. Changes in serotoninergic receptors 1A and 2A in the piglet brainstem after intermittent hypercapnic hypoxia (IHH) and nicotine. Brain research. 2007;1152:17–26.

19. Meyerholz DK, Ofori-Amanfo GK, Leidinger MR, et al. Immunohistochemical Markers for Prospective Studies in Neurofibromatosis-1 Porcine Models. J Histochem Cytochem. 2017;65(10):607–618.

20. Steen RG, Taylor JS, Langston JW, et al. Prospective evaluation of the brain in asymptomatic children with neurofibromatosis type 1: relationship of macrocephaly to T1 relaxation changes and structural brain abnormalities. AJNR. American journal of neuroradiology. 2001;22(5):810–817.

21. Cutting LE, Cooper KL, Koth CW, et al. Megalencephaly in NF1: predominantly white matter contribution and mitigation by ADHD. Neurology. 2002;59(9):1388–1394.

22. Duffner PK, Cohen ME, Seidel FG, Shucard DW. The significance of MRI abnormalities in children with neurofibromatosis. Neurology. 1989;39(3):373–378.

23. Costa RM, Federov NB, Kogan JH, et al. Mechanism for the learning deficits in a mouse model of neurofibromatosis type 1. Nature. 2002;415(6871):526–530.

24. Balestrazzi P, de Gressi S, Donadio A, Lenzini S. Periaqueductal gliosis causing hydrocephalus in a patient with neurofibromatosis type 1. Neurofibromatosis. 1989;2(5-6):322–325.

25. Okon EA, Ikpeme EV, Udensi OU, et al. Computational Analysis of Evolutionary Relationship of a Family of Cold Shock Proteins in Ten Mammalian Species. Journal of Advances in Biology & Biotechnology. 2017;16(2):1-14.

26. Gene Expression Database (GXD). http://www.informatics.jax.org. Accessed November 28, 2018.

27. Wong AM, Patel NV, Patel NK, et al. Macrosialin increases during normal brain aging are attenuated by caloric restriction. Neuroscience letters. 2005;390(2):76–80.

28. Ligon KL, Kesari S, Kitada M, et al. Development of NG2 neural progenitor cells requires Olig gene function. Proceedings of the National Academy of Sciences of the United States of America. 2006;103(20):7853–7858.

29. Cammer W, Zhang H. Atypical glial cells in demyelinated and hypomyelinated mouse brains. Brain research. 1999;837(1-2):188–192.

30. Mouse brain tissue atlas. https://www.proteinatlas.org/ENSG00000180176-TH/tissue/mouse+brain. Accessed November 28, 2018.

31. Boccazzi M, Ceruti S. Where do you come from and what are you going to become, reactive astrocyte? Stem cell investigation. 2016;3:15.

32. Uhlén M, Fagerberg L, Hallström BM, et al. Tissue-based map of the human proteome. Science. 2015;347(6220):1260419.

33. Dean JM, Farrag D, Zahkouk SA, et al. Cerebellar white matter injury following systemic endotoxemia in preterm fetal sheep. Neuroscience. 2009;160(3):606–615.

34. Shitaka Y, Tran HT, Bennett RE, et al. Repetitive closed-skull traumatic brain injury in mice causes persistent multifocal axonal injury and microglial reactivity. Journal of neuropathology and experimental neurology. 2011;70(7):551–567.

35. Hart AD, Wyttenbach A, Perry VH, Teeling JL. Age related changes in microglial phenotype vary between CNS regions: grey versus white matter differences. Brain, behavior, and immunity. 2012;26(5):754–765.

36. Galvao RP, Kasina A, McNeill RS, et al. Transformation of quiescent adult oligodendrocyte precursor cells into malignant glioma through a multistep reactivation process. Proceedings of the National Academy of Sciences of the United States of America. 2014;111(40):E4214–4223.

37. Parri HR, Gould TM, Crunelli V. Sensory and cortical activation of distinct glial cell subtypes in the somatosensory thalamus of young rats. The European journal of neuroscience. 2010;32(1):29–40.

38. Reynolds R, Hardy R. Oligodendroglial progenitors labeled with the O4 antibody persist in the adult rat cerebral cortex in vivo. Journal of neuroscience research. 1997;47(5):455–470.

39. Levine JM, Stincone F, Lee YS. Development and differentiation of glial precursor cells in the rat cerebellum. Glia. 1993;7(4):307–321.

40. Castro-Caldas M, Neves Carvalho A, Peixeiro I, Rodrigues E, Lechner MC, Gama MJ. GSTpi expression in MPTP-induced dopaminergic neurodegeneration of C57BL/6 mouse midbrain and striatum. Journal of molecular neuroscience: MN. 2009;38(2):114–127.

41. Robinson AP, Rodgers JM, Goings GE, Miller SD. Characterization of oligodendroglial populations in mouse demyelinating disease using flow cytometry: clues for MS pathogenesis. PloS one. 2014;9(9):e107649.

42. Sharaf A, Rahhal B, Spittau B, Roussa E. Localization of reelin signaling pathway components in murine midbrain and striatum. Cell and tissue research. 2015;359(2):393–407.

43. Diggs-Andrews KA, Tokuda K, Izumi Y, Zorumski CF, Wozniak DF, Gutmann DH. Dopamine deficiency underlies learning deficits in neurofibromatosis-1 mice. Annals of neurology. 2013;73(2):309–315.

44. Murphy KP, Carter RJ, Lione LA, et al. Abnormal synaptic plasticity and impaired spatial cognition in mice transgenic for exon 1 of the human Huntington’s disease mutation. The Journal of neuroscience: the official journal of the Society for Neuroscience. 2000;20(13):5115–5123.

45. Manohar S, Paolone NA, Bleichfeld M, Hayes SH, Salvi RJ, Baizer JS. Expression of doublecortin, a neuronal migration protein, in unipolar brush cells of the vestibulocerebellum and dorsal cochlear nucleus of the adult rat. Neuroscience. 2012;202:169–183.

46. Zhao C, Eisinger B, Gammie SC. Characterization of GABAergic neurons in the mouse lateral septum: a double fluorescence in situ hybridization and immunohistochemical study using tyramide signal amplification. PloS one. 2013;8(8):e73750.

47. Nelson TE, King JS, Bishop GA. Distribution of tyrosine hydroxylase-immunoreactive afferents to the cerebellum differs between species. The Journal of comparative neurology. 1997;379(3):443–454.

48. Asmus SE, Anderson EK, Ball MW, et al. Neurochemical characterization of tyrosine hydroxylase- immunoreactive interneurons in the developing rat cerebral cortex. Brain research. 2008;1222:95–105.

49. Borsook D. Neurological diseases and pain. Brain: a journal of neurology. 2012;135(Pt 2):320–344.

50. Yu D, Liu F, Liu M, et al. The inhibition of subchondral bone lesions significantly reversed the weight-bearing deficit and the overexpression of CGRP in DRG neurons, GFAP and Iba-1 in the spinal dorsal horn in the monosodium iodoacetate induced model of osteoarthritis pain. PloS one. 2013;8(10):e77824.

51. Rice FL, Xie JY, Albrecht PJ, et al. Anatomy and immunochemical characterization of the non- arterial peptidergic diffuse dural innervation of the rat and Rhesus monkey: Implications for functional regulation and treatment in migraine. Cephalalgia: an international journal of headache. 2017;37(14):1350–1372.

52. Tognatta R, Miller RH. Contribution of the oligodendrocyte lineage to CNS repair and neurodegenerative pathologies. Neuropharmacology. 2016;110(Pt B1):539-547.

53. Arnold SE, Han LY, Clark CM, Grossman M, Trojanowski JQ. Quantitative neurohistological features of frontotemporal degeneration. Neurobiology of aging. 2000;21(6):913–919.

54. Nielsen HM, Ek D, Avdic U, et al. NG2 cells, a new trail for Alzheimer’s disease mechanisms? Acta neuropathologica communications. 2013;1:7.

55. Gourfinkel-An I, Parain K, Hartmann A, et al. Changes in GAD67 mRNA expression evidenced by in situ hybridization in the brain of R6/2 transgenic mice. Journal of neurochemistry. 2003;86(6):1369–1378.

